# Intrinsic Mechanics of Human Stem Cell Derived Aortic Smooth Muscle Cells Support a Developmental Basis for Aneurysm Localization in Marfan Syndrome

**DOI:** 10.1101/2023.10.03.560723

**Authors:** Robert J. Wiener, Helen Orins, Kevin D. Costa

## Abstract

Marfan Syndrome (MFS), a connective tissue disorder caused by a mutation in the fibrillin-1 gene, occurs in approximately 1 in 5,000 people worldwide. As an important constituent of the extracellular matrix, mutated fibrillin-1 in Marfan Syndrome leads to aortic medial degeneration, aneurysm, and dissection. TGFβ in the matrix, which is controlled by fibrillin-1, is known to cause pathological effects in smooth muscle cells (SMCs) within the aortic wall during MFS. TGFβ as well as other cytokines have been shown to impact neural crest derived SMCs differently than mesodermal derived SMCs. Furthermore, outcomes of variable cytokine responsiveness of neural crest SMCs are compounded by genetically imposed changes to neural crest SMC integrin distributions in MFS. Thus, it has been hypothesized that neural crest derived SMCs, which give rise to ascending aortic SMCs, are intrinsically mechanically susceptible to aneurysm formation in MFS. This hypothesis has been linked to the clinical observation of aneurysm formation preferentially occurring in the ascending versus descending aorta in MFS. We aim to test the hypothesis that aortic smooth muscle cells (ASMCs) have intrinsic mechanobiological properties which cause cell weakening in Marfan Syndrome. Human induced pluripotent stem cells (hiPSC) from Marfan patients and healthy volunteers were differentiated into either ascending- or descending-ASMCs via their respective developmental lineages, and cultured to either an early (6 days) or late (30 days) stage of post-differentiation maturation. Mass spectrometry-based proteomics of early-stage iPSC-ASMCs revealed an array of depleted proteins unique to MFS ascending-SMCs that were associated with cell mechanics and aortic aneurysm. Targeted examination of the proteomics dataset revealed intracellular proteins (ACTA2, CNN1, TAGLN) were significantly depleted in MFS ascending-ASMCs. The intrinsic, matrix-independent, hiPSC-ASMC stiffness quantified by atomic force microscopy (AFM) revealed that MFS ascending-ASMCs, but not descending-ASMCs, were significantly less stiff than healthy, at the late cell-maturation stage (p<0.0005). Late-stage ascending- and descending-ASMCs also showed clear functional impairments via calcium flux in MFS. AFM revealed a similar mechanical phenotype in early-stage ASMCs, with MFS ascending-ASMCs, but not descending-ASMCs, being significantly less stiff than healthy (p<0.005). In summary, this study supports an emerging hypothesis of ontogenetic predisposition for aneurysm susceptibility in Marfan Syndrome based on locally altered mechanobiology of developmental origin-specific ASMC subtypes. This may lead to new cell-targeted approaches for treating aortic aneurysm in patients with MFS.

## Introduction

Marfan Syndrome (MFS) is a heritable autosomal dominant disease, caused by a mutation in the fibrillin-1 gene, that manifests as degenerative connective tissues across multiple organ systems. Marfan Syndrome often causes ectopia lentis (dislocation of the eye lens), scoliosis, and mitral valve prolapse. Almost synonymous with Marfan Syndrome is aortic aneurysm, which is also its most deadly manifestation. Progressive aortic wall degeneration, aneurysm, and aortic dissection predominantly in the ascending aorta (Stanford Type A) have been widely attributed to locally elevated hemodynamics, with MFS deemed as a matrix disease ^1,2^. However, it is also known that ascending aortic smooth muscle cells (ASMCs) come from a distinct embryologic developmental origin ^3–6^. While ascending-ASMCs are ectodermal derived, descending and other ASMCs are mesodermal derived, with each SMC sub-type occupying a distinct region within the aorta ^3–5,7–9^. While investigation of the origin-specific differences in ASMCs can improve our understanding of their role in MFS, dissecting their involvement has been challenging with animal models in vivo. This has motivated the use of human-specific model systems based on patient-derived pluripotent stem cells and lineage-specific differentiation of smooth muscle cells, as described in recent published work ^10–12^.

Despite promising therapeutic results in rodent models of MFS ^13–17^, Losartan has underperformed in clinical trials ^18,19^. One theory is that preventing aneurysm formation caused by aberrant TGFβ signaling is dependent on age. Researchers found that neutralization of TGFβ before postnatal day 16 (P16) in the *Fbn1*^*mgR/mgR*^ Marfan mouse exacerbated aneurysm formation, while neutralization after P45 mitigated aneurysm formation ^16^. While TGFβ involvement in MFS is clear, the source of aortic weakening and aneurysm is not. Some have suggested that it is endothelial NOS activation which drives SMC dysfunction, but structural and mechanical studies have shown that aneurysm in *Fbn1*^*mgR/mgR*^ MFS mice causes weakening primarily in the media layer of the aortic wall, not the intima ^13^. Therefore, it makes sense to target the mechanics of media-residing ASMCs in aortic aneurysm ^20,21^.

Recently a new ASMC-based theory has begun to gain traction. While Marfan Syndrome patients experience matrix degeneration throughout their entire aorta (both ascending and descending), dissection most often occurs in the ascending aorta region (Type A), with a prevalence of about 80% compared to descending (Type B) dissection; this suggests that matrix degeneration could be necessary but not sufficient for aortic aneurysm and dissection in MFS. Another active tissue component may be a key driver; aortic smooth muscle cells. While ASMCs have been studied in the context of MFS, they are often obtained from patient biopsies, and are therefore already diseased and exposed to matrix in an advanced pathological state. Novel protocols have been published providing a way to generate aortic region-specific smooth muscle cells from human iPSCs to study diseases such as MFS ^10,11,22^. Using a modified version of the Cheung et al., 2014 protocol researchers discovered that not all ASMCs behave the same in MFS; lateral plate mesoderm-derived ASMCs, in comparison to neural crest-derived ASMCs, had increased integrin αV and reduced mannose receptor C-type 2, but exhibited reduced adhesion to Collagen I & IV ^23^. Interestingly, a team led by Dr. Sinha at the University of Cambridge found that late-stage neural crest derived (ascending) ASMCs had increased integrin β1 in MFS, while mesodermal ASMCs did not, and that the associated increase in downstream Itgb1 signaling (P-p38, KLF4) was also present in primary human aortic tissue from MFS patients ^12^. Moreover, they showed that CRISPR correction of the MFS mutation in neural crest SMCs corrected the fibrillin-1 phenotype, and ameliorated the increase in mechanobiologically associated MMP10 levels. These findings suggest that differentiation of iPSC to region-specific ASMCs provides a translatable model for studying MFS, and furthermore allow researchers to dissect the roles of cells vs. matrix in mechanobiologically driven aortic pathologies. This study aims to elucidate, in a translatable human iPSC-based model system, the intrinsic mechanobiological mechanisms unique to neural crest derived ASMCs in MFS. We also aim to connect the ASMC mechanical phenotype in 2-dimensions to their behavior in a 3-dimensional environment for a better understanding of regional aneurysm susceptibility in Marfan Syndrome.

## Methods

### Patient Cell Lines

As part of the NIH-Common Fund Library of Integrated Network-Based Cellular Signatures (LINCS) program ^24^ an array of 40 healthy individuals (based on rigorous clinical evaluations) were selected to create hiPSCs by reprogramming the patient’s dermal fibroblasts via mRNA reprogramming kit (Stemgent, Cat # 00-0071) in combination with the microRNA Booster Kit (Stemgent, Cat# 00-0073) and/or the CytoTune-iPS 2.0 Sendai Reprogramming Kit (Thermo Fisher Scientific, Cat# A16517) ^25^.

Through the LINCS project, the derived hiPSCs underwent cytogenetic reports, short tandem repeat authentication, pluripotency analysis, whole-genome sequencing, genomic ancestry determination, and Mendelian disease gene and risk assessment. Healthy hiPSC lines MSN09 (25-year-old, white female) and MSN22 (26-year-old, white female) were selected as controls for this study. Marfan iPSC lines were obtained from the Coriell Institute for Medical Research (Camden, NJ). The hiPSC line MFS44 donor (7-year-old female, Coriell_ID: GM21944) had a heterozygous mutation in FBN1, 3976T>C (FBN1p.C1326R), experienced ascending aortic aneurysm, and surgical replacement of the aortic root. The MFS60 cell line donor (28-year-old female, Coriell_ID: GM21960) had a heterozygous mutation in FBN1 is 4082G>A (FBN1p.C1361Y), experienced ascending aortic aneurysm, dissection, and surgical replacement of the aortic root. All Marfan iPSC lines were also derived from dermal fibroblasts. Note that all ages mentioned were the age of the donor at biopsy.

### HiPSC Culturing and Lineage-Specific ASMC Differentiations

HiPSCs were grown on flat-bottom plastic 6-well tissue culture plates coated with hESC-qualified Matrigel (Corning, cat# 354277). HiPSCs were maintained in a 37°C incubator at 5% CO_2_ and 95% humidity using StemFlex media (Thermo Cat#: A3349401), and replaced with fresh media (2mL per well) every other day. To passage, cells were treated at room temperature with ReLeSR (Stem Cell Technologies, Cat#: 05872) for 1 minute followed by 5 minutes at 37°C, then dissociated using StemFlex with Thiazovivin (1:5000), and plated on new Matrigel coated 6-well plates at a ratio of 1:6. After 24 hrs, plated cells were given fresh StemFlex without Thiazovivin. A detailed protocol for passaging and cryopreservation in a defined media can be found in ^26^. A detailed protocol for differentiation of hiPSCs into lineage-specific aortic smooth muscle cells can be found in Cheung et al., 2014 Nature Protocols ^10^. For this study, iPSCs were grown to about 40% confluence in StemFlex, then switched to a chemically defined medium. Note that differentiations took place at 37°C, 5% CO_2_ normoxic conditions. Chemically defined medium (CDM) is made by combining 250 mL of Iscove’s Modified Dulbecco’s Medium (ThermoFisher, Cat#: 12440053), 250 mL of Ham’s F-12 Nutrient Mix (ThermoFisher, Cat#: 11765054), 5 mL Chemically Defined Lipid Concentrate (ThermoFisher, Cat#: 11905031), 250 μL Transferrin (30 mg/mL, Roche, Cat#: 652202), 20 μL of 1-Thioglycerol (Sigma, Cat#: M6145), 5 mL penicillin-streptomycin, then vacuum filtering, then adding 350 μL of sterile Insulin (10 mg/mL, Roche, Cat#: 1376497). After being switched to CDM, hiPSCs were primed for induction with 12 ng/mL FGF2 (R&D Systems, Cat# 233-FB) and 10 ng/mL activin A (R&D Systems, cat# 338-AC) for 24 hours. For induction of intermediate developmental cell populations, different cell fates were specified by treatment with varying growth factors for each lineage following Cheung et al. ^10^. Neural crest intermediates were defined with 7 days of 12 ng/mL FGF2 and 10 μM of the activin–nodal inhibitor SB431542 (Sigma, cat# S4317) in CDM, while paraxial mesoderm intermediates were defined with 24 hrs of 20 ng/mL FGF2, 10 μM LY294002 (Sigma, Cat# L9908) a PI3K inhibitor, and 10 ng/mL BMP4 (R&D Systems, Cat#: 314-BP), followed by 4 more days of FGF2 and LY294002 in CDM. To confirm positive intermediate populations, neural crest derivatives were stained for positive expression of Nestin, and paraxial mesoderm derivatives were stained for positive TBX18 expression. Once intermediate populations were defined, cells were passaged to new Matrigel coated culture plates for terminal differentiations. This was done by dissociating cells for 3 min at 37°C using TrypLE Express (ThermoFisher, Cat#: 12605010), neutralizing with 1:10 volumes (CDM), spinning down for 3 min at 250 g, resuspending in CDM with 2 ng/mL TGFβ1 and 10 ng/mL PDGF-BB, then replating at 30,000 cells per cm^2^. After 24 hrs the media (CDM+TGFβ1+PDGF-BB) was replaced. Terminal aortic smooth muscle cell differentiation occurred for 12 days, changing media every other day with 2 ng/mL TGFβ1 (PeproTech, Cat# 100-21C) and 10 ng/mL PDGF-BB (PeproTech, Cat# 100-14B) in CDM. During this time, differentiating cells would be passaged to avoid overgrown cell clumps, or over 95% confluent. Once differentiation was complete on day 13, ASMCs were switched to Advanced Dulbecco’s Modified Eagle Medium/Ham’s F-12 (ADMEM) (ThermoFisher, Cat# 12634010) supplemented with 10% Fetal Bovine Serum (FBS) and allowed to mature for either 6 days (early-stage) or 30 days (late-stage) ^10–12^. To replate iPSC-ASMCs for experiments, cells were again released with TrypLE Express, neutralized in 10 parts media, spun down at 250g for 3 min, then plated at 20,000 cells/cm^2^ in ADMEM+FBS supplemented with an additional 10% of conditioned cell media. Importantly for experimentation, ASMCs were plated on preconditioned gelatin plates. Preconditioned gelatin plates were made by coating 50mm cover-glass bottom Petri dishes (WPI Inc, Cat#: FD5040) with 0.1% gelatin (Sigma, Cat#: G1890) for 1 hr at room temperature, then washing and replacing with ADMEM+FBS to condition overnight at 37°C.

### Primary Human Aortic Smooth Muscle Cell Culture

Primary human aortic smooth muscle cells (HAOSMCs) from Cell Applications (healthy 26-year-old, white male, Cat#: 354-05a) were grown on tissue culture treated Petri dishes (Corning, Cat#: 430166) using complete HAOSMC growth media (Cell Applications, Cat#: 311-500). After one passage, the cells were adapted to Advanced Dulbecco’s Modified Eagle Medium/Ham’s F-12 (ADMEM) (ThermoFisher, Cat# 12634010) supplemented with 10% Fetal Bovine Serum (FBS) in order to match iPSC-ASMC culture conditions. In plating for experiments, HAOSMCs were dissociated with Trypsin-EDTA (0.25%) for 3 minutes, neutralized in 10 parts media, spun down at 250g for 3 min and then plated at 20,000 cells/cm^2^ on 0.1% gelatin coated dishes. Experiments were performed on cells below passage 7.

### Live iPSC-ASMC Calcium Signaling

ASMCs were plated on a gelatin coated preconditioned (see 4.3.2) 35 mm high glass bottom μ-Dishes (ibidi, Cat#: 81158) using ADMEM+10%FBS and allowed to reach at least 80% confluency. Cells were then serum starved (switched to only ADMEM) for 48 hrs prior to experiment. On the day of experiment, cells were loaded in loading media (DMEM+glutamine without phenol red (ThermoFisher, Cat#: 21041025)) containing 2.5 μL/mL of Fluo-4AM (ThermoFisher, Cat#: F14201) for 1 hr at 37°C, then washed with loading media for 20 min at 37°C. Cells were then switched into imaging media (439 mL dH_2_O, 50 mL 10x HBSS, 5 mL 10x sodium pyruvate, 5 mL HEPES, 460 μL 1M MgCl_2_, 450 μL 1M CaCl_2_) and placed on an Olympus IX-81 inverted microscope with heated stage (set to 37°C) and active vibrational isolation. Fluorescence was controlled via a Lambda LS Sutter instrument and captured using a Hamamatsu EM-CCD camera. All experimental parameters were prescribed using Olympus Metamorph Basic software. First at x10 magnification a baseline cell fluorescence (GFP) was captured for 30 sec, then an equal part of imaging media was added and captured for another 30 sec (as vehicle control); finally, 2.5 mM of acetylcholine receptor agonist, Carbachol (Abcam, Cat#: ab141354) was added and recorded for the remaining 240 sec. Note that carbachol was added such that the final volume of fluid on the cells yielded a final concentration of 2.5 mM. The experiment images were acquired via the GFP channel at 250 ms exposure time, each frame was the result of 3 GFP acquisitions averaged, which yielded 1 frame/sec. This experiment took place over 300 second’s total. All images were acquired using no autofocus. After recording was complete, cellular calcium (fluorescence) fluctuations in response to carbachol was quantified in FIJI (ImageJ, fiji.sc). All 300 frames were imported as an image sequence into FIJI, next a background area and fluorescent cell area were selected. For each of the 300 frames, the average fluorescence intensity was calculated (both for the background and cell). To acquire the relative change in fluorescence ΔF, defined by:

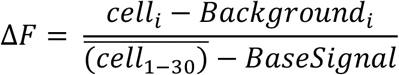

where *i* is frame 1 to 300, and

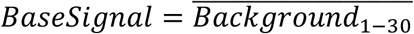

Note frames 1-30 for *BaseSignal* use the “background” selected area, and *cell* use the “cell” selected area. Frames 31-59 are used to evaluate the vehicle effect. The overhead bar indicates an average of frames. Each cell was analyzed individually and the collective results were averaged for group metrics.

### Live ASMC Mechanics Measured via Atomic Force Microscopy

ASMCs were plated at a low density to visualize cell morphology and to allow isolated cell measurements (see 4.3.2 and 4.3.3). Each dish of cells was secured to the stage of an MFP-3D-BIO (Asylum Research Inc) Atomic Force Microscope (AFM) which was equipped with environmental vibration isolation and temperature control (set to 37°C). An AFM probe (Asylum Research Inc, Cat#: TR400PB) was used to perform nano-indentation measurements on individual cells. The TR400PB contains a silicon nitride, gold-alloy coated, triangular cantilever with a nominal spring constant of 0.09 N/m, and a pyramidal-shaped tip (nominal radius 40 nm). This probe has features well suited for micromechanical measurements of live cells ^27,28^. The AFM was controlled by Igor Pro 6.3 software (Wavemetric Inc). First a cytoplasmic area intermediate between the cell edge and nucleus was identified. The AFM probe was then prescribed to indent the cell body at a rate of 5 μm/sec until it reached a cantilever deflection (trigger) of 20 nm, this equated to a contact force of approximately 3 nN. To account for intracellular heterogeneity, an array of 16 indentations over a 5×5-μm square region was performed. The probe’s force-distance response (force curve) from each cell indentation was fit using a Hertzian model to the advance portion of the force curve (10-90% post cell contact) which yielded a local elastic modulus ^29,30^. Individual indentations were inspected, and clearly misfit curves were eliminated. If over 50% of the 16 indentations passed, that array was accepted. The median of all remaining elastic moduli for the array of 16 indentations was taken as the apparent elastic modulus of the cell ^29,30^.

### Mass-Spectrometry Based Quantification of Proteins

Samples representing both developmental origin (neural crest vs. paraxial mesoderm) and genotype (WT vs. MFS) were prepared for Tandem Mass Tags (TMT) based shotgun proteomics. Briefly, healthy iPSCs (MSN09) and Marfan iPSCs (MFS60) were each differentiated to both ascending and descending ASMCs, then harvested at early-stage maturation (6 days in serum post terminal ASMC differentiation). Four differentiations were used for each group, yielding 16 total samples (4 conditions, each with 4 biological replicates). To harvest, cells were first thoroughly washed in PBS (two times with 2mL per well) to remove any serum, then left in ice cold PBS for 8 min to dissociate. Next, cell dissociation was completed by pipette scraping manually in PBS and spun down for 3 min at 300g. Finally, the cell pellet was resuspended in 1 mL of DPBS with HALT protease inhibitor (ThermoFisher, Cat#: 78430) and again spun down for 3 min between 300-1000g. The supernatant was removed and the sample was flash frozen in liquid nitrogen, then stored at -80°C. Note that Marfan ascending ASMCs seemed to be the most delicate sample and were spun down at 300g while the other samples were spun down at 1000g.

Samples were then sent to the Proteomics Research Facility at the University of Rutgers, Newark NJ and quantified using the following methods. Each sample was lysed and sonicated in a lysis buffer [8 M urea, 100 mM TEAB (pH 8.5)] that contained protease inhibitor cocktail, and Phosphatase Inhibitor Cocktail I & II (Sigma). The samples were moved to fresh tubes and 100 ug of each sample (confirmed by BCA protein assay) was collected for processing. Sample proteins were reduced with dithiothreitol (DTT), and subsequently, free thiols were blocked by iodoacetamide (IAM). First, proteins were digested with Lys-C for 4 hours at 37 °C, then diluted to a final urea concentration of 1 M. Next, digestion was performed with the addition of 4 ug of trypsin, and incubated at 37 °C overnight. The resulting peptides derived for each sample were labeled with one of the 16 plex TMT tags as per the manufacturer’s instructions (ThermoFisher). The labeled peptides were combined, dried in a SpeedVac, and desalted by C_18_ cartridges (Thermo Scientific). Finally, the combined TMT-labelled sample was fractionated using a high pH RPLC (Waters XBridge BEH C_18_ column, 5μm, 4.6×250mm). Sixty fractions were collected and combined into 13 fractions; each fraction was dried using a Speedvac and desalted using C_18_ cartridges

Peptides from the 13 fractions were analyzed *via* a low pH RPLC-MS/MS using a Fusion Lumos Orbitrap Mass spectrometer coupled with an UntiMate 3000 RSLC nano system (Thermo Scientific). Briefly, the peptides were directly loaded onto a reversed phase column (Acclaim PepMap C_18_ column, 75 μm × 50 cm, 2 μm, 100 Å). The peptides were eluted using a 3-hr binary gradient of solvent A (2% acetonitrile (ACN) in 0.1% formic acid (FA)) and solvent B (85% ACN in 0.1% FA) at a flow rate of 300 nL/min. The eluted peptides were introduced into a nano electrospray source on an Orbitrap Fusion Lumos MS instrument, with a spray voltage of 2 kV and a capillary temperature of 275 °C. The MS spectra were acquired in a positive ion mode with a resolution of 120,000 full-width at half maximum (FWHM). The MS scan range was between m/z 350 to 1600. The Higher Energy Collision Dissociation (HCD) MS/MS spectra were acquired in a data-dependent manner with a quadrupole isolation window of 0.7 m/z. The normalized HCD collision energy was set to 34%. The peptide fragments were detected in the Orbitrap analyzer at a resolution of 50,000, with the AGC target set to a normalized target of 150%. The dynamic exclusion was set to 60 sec.

The MS/MS spectra were searched against an UniProt human database (downloaded on 10/28/2021, 77,897 human protein sequences) using the Sequest search engines within the Proteome Discoverer platform (version 2.4, Thermo Scientific). The following search parameters were used: (i) fixed modifications including TMT 16plex (K), TMT 16plex (N-terminal), and Carbamidomethylation (C); (ii) variable modifications included oxidation (M); (iii) trypsin was selected as the digestive enzyme; (iv) up to two missed cleavage sites were allowed; (v) the peptide precursor mass tolerance was 10 ppm; and MS/MS mass tolerance was 0.1 Da. A false discovery rate (FDR) was maintained at under 1% based on the filter in the Percolator module. Protein quantitation was normalized based on the total amounts of peptides including both unique and razor peptides. In kind methods regarding protein extraction, labeling, LC-MS/MS, and protein identification can be found here ^31–33^.

To compare protein abundances between groups and evaluate for differentially abundant proteins (DAPs) the R programming language, Linear Models for Microarray Data (LIMMA) package (Bioconductor.org)^34^ was used to calculate log_2_ fold-change and adjusted p-value to control the false discovery date ^35^. The first two DAP datasets were based on comparisons within origin (i.e., ascending WT/MFS and descending WT/MFS). The final DAP dataset was the ascending set filtered for any overlap with descending, this returned DAPs uniquely different in ascending MFS versus WT. The final subset of data considered were the most significant DAPs in ascending MFS which were selected based on the TREAT method ^36^, using a log_2_ fold-change cutoff of ±1 and adjusted p-value of 10^-4^. The resulting arrays of DAPs were fed into Enrichr suite ^37–39^ (https://maayanlab.cloud/Enrichr/) to identify highly enriched biological processes implicated in the disease phenotype.

### Immunofluorescence

Samples were fixed in 4% paraformaldehyde (PFA) in PBS at room temperature (10 min for cells) and were then permeabilized with 0.1% Triton-X (5 min for cells) and thoroughly washed in PBS. Samples were then blocked with goat serum (1 hr at 37°C for cells). To stain the samples, an antibody mixture containing 1-part DAKO (Agilent, Cat#: S080983-2) and 4-parts goat serum was mixed to create a desired antibody dilution:

| Primary antibody dilution table |  |  |
| --- | --- | --- |
| Primary antibody | Abbreviation | Dilution |
| α-smooth muscle actin (abcam, Cat#: ab124964) | αSMA | 1:300 |
| CNN1 (sigma, Cat#: C2687) | Calponin-1 | 1:500 |
| TAGLN (abcam, Cat#: ab14106) | Transgelin | 1:500 |
| T-box transcription factor 18 (abcam, Cat#: ab115262) | TBX18 | 1:200 |
| Nestin, FITC (abcam, Cat#: ab187846) | Nestin | 1:200 |
|  |  | **Cell** |

Secondary antibodies were made to the same primary dilution but incubated with samples for 1 hr at 37°C.

| Secondary antibodies |
| --- |
| Goat anti-Rabbit IgG, AF488 (ThermoFisher Cat#: A-11008) |
| Donkey anti-Rabbit IgG, AF546 (ThermoFisher Cat#: A10040) |
| Goat anti-Mouse IgG, Cyanine3 (ThermoFisher Cat#: A10521) |

## Results

### Underrepresented Proteins Unique to Marfan Ascending ASMCs Offer Unique Insight into the Early-stage Disease Phenotype

We evaluated our TMT proteomics dataset for differentially abundant proteins unique to Marfan ascending ASMCs. Two differentially abundant protein (DAP) datasets were created comparing genotypes within each origin (i.e. ascending Healthy/MFS and descending Healthy/MFS), the final dataset was created by filtering out any DAPs in the ascending group that overlapped with the descending. This final dataset provided DAPs uniquely depleted or abundant in ascending Marfan ASMCs. **Figure 1a** shows the data from the final dataset as a volcano plot with p-value significance on the y-axis and fold-change on the x-axis. In selecting the most significant targets of this dataset we captured depleted proteins with an adjusted p-value greater than 10^-4^ and a log_2_ fold-change less than - 1, which yielded 88 significantly depleted proteins. Similarly, we captured abundant proteins with an adjusted p-value greater than 10^-4^ and a log_2_ fold-change greater than 1, which yielded 148 significantly abundant proteins. After conducting an enrichment analysis using the top depleted proteins we identify key biological processes linked to that subset of proteins (**Figure 1b,c,d**). This enrichment suggested that our significantly depleted proteins were implicated with the biological processes of disrupted focal adhesion and cytoskeletal regulation (**Figure 1b**), gene ontology of extracellular matrix organization (**Figure 1c**), and clinical outcome of thoracic aortic dissection (**Figure 1d**).

**Figure 1:**
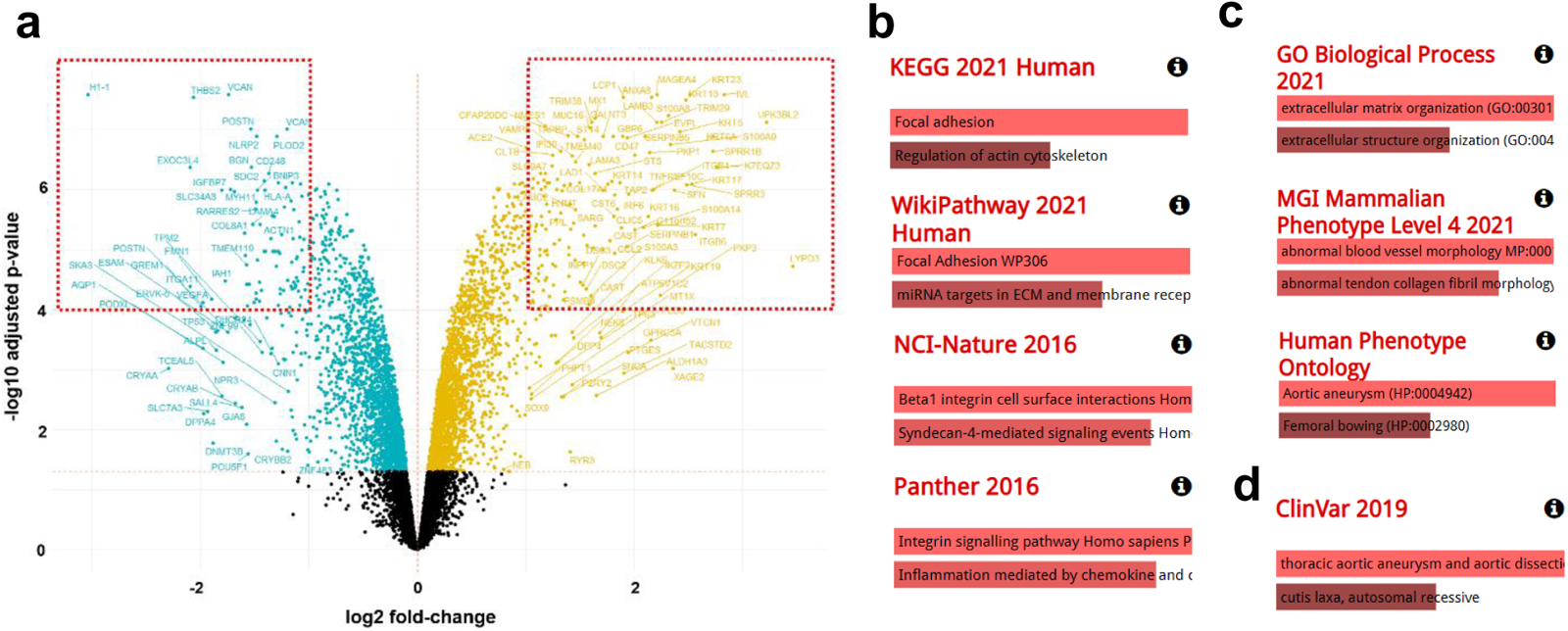
Differential abundance analysis reveals unexplored mechanobiological proteins uniquely depleted in ascending MFS ASMCs. (a) Volcano plot of differentially abundant proteins unique to ascending ASMCs, where proteins significantly depleted in MFS ascending ASMCs are blue, and significantly abundant in MFS ascending ASMCs are yellow. Significance was defined by having an adjusted p-value of <10^-4 and log2 fold-change beyond ±1.0. Enrichr software shows enrichment of significantly depleted protein subset (b) biological process, (c) gene ontology, and (d) disease process.

### Proteins Associated with ASMC Mechanics are Depleted in Marfan Ascending Aortic Cells

By exploring the TMT proteomic dataset in a targeted manner, we examined the proteins levels of key regulators identified in the enrichment analysis of (**Figure 1**). The abundance of smooth muscle specific proteins commonly associated with cell mechanics ^29,30,40^ were significantly depleted in Marfan vs. healthy ascending ASMCs (α-smooth muscle actin (ACTA2), and calponin-1 (CNN1), and transgelin (TAGLN)) (**Figure 2a**). Furthermore, Marfan ascending ASMCs has depleted proteins essential for the cell’s contractile machinery (myosin heavy chain-11 (MYH11), and myosin light chain-6B, (MYL6B)) **Figure 2b**). Importantly, proteins commonly used as controls were found to be not significantly different across ASMC groups (beta actin (ACTB) and tubulin beta 1 (TUBB1)), supporting consistency of the technical preparations (**Figure 2c**). It is interesting to note that while statistical significance was observed for the ascending ASMC groups, this was not so for the descending ASMC groups. This could partly reflect the larger variability in descending samples compared to the ascending samples.

**Figure 2:**
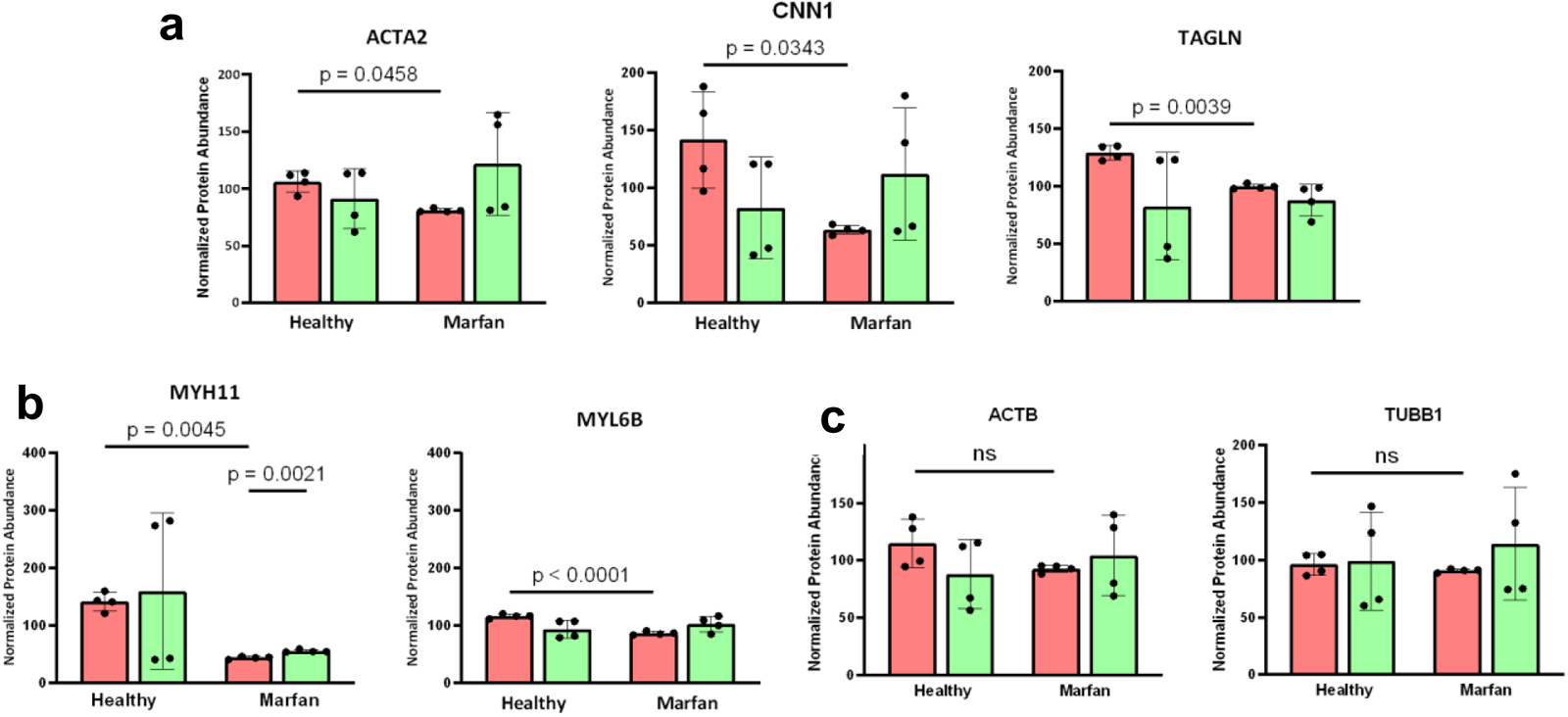
Early-stage ascending ASMCs in Marfan Syndrome have depleted proteins related to the disease phenotype. (a) ASMC cytoskeletal and stiffness related proteins; ACTA2 (smooth muscle actin), CNN1 (calponin), TAGLN (transgelin). (b) contractile ASMC proteins; MYH11 (myosin heavy chain-11), MYL6B (myosin light chain-6B). (c) typical control proteins; ACTB (beta actin), TUBB1 (beta tubulin). All ASMCs were harvested at an early stage from either cell line MSN09 or MFS44. Bars represent mean±SD. p-values were calculated using an unbalanced multi-way ANOVA with Tukey-Kramer post hoc

### AFM Reveals Late-stage Ascending ASMCs are Uniquely Compliant in Marfan Syndrome

Using Atomic Force Microscopy (AFM), we measured the apparent cell stiffness of ascending and descending ASMCs, at a late cell-maturation stage. **Figure 3** presents the intrinsic elastic modulus of region-specific iPSC-ASMCs from different genotypes subject to the same environment. AFM revealed that in healthy late-stage ASMCs, ascending cells tended to be stiffer than descending. Conversely in Marfan, ascending tended to be less stiff (more compliant) than descending. Importantly, Marfan ascending-ASMCs were the least stiff compared to both healthy ascending (p<0.0001) and healthy descending (p=0.0010) (**Fig. 3a**). Interestingly, while Marfan ascending-ASMCs were significantly less stiff than both healthy ASMC groups, Marfan descending ASMCs were only significantly less stiff than healthy ascending-ASMCs (p<0.0001) (**Fig. 3a**). **Figure 3b** shows that the AFM measured stiffness of primary human ASMCs from a healthy adult is within range of our healthy iPSC-derived ASMCs. Additionally, all iPSC-dervied and primary ASMCs showed positive staining for smooth muscle cell markers (**Fig. 3d**). While positive for both cytoskeletal (αSMA, SM22α) and calcium signaling (CNN1) related protiens, their relative abundance was not yet quantified. Taken together, these data show that SMC-positive iPSC-derived ASMCs have *in vitro* stiffness comparable to primary human ASMCs, and importantly that Marfan iPSC-derived ascending-ASMCs are intrinsically less stiff than healthy. This suggests that the genetics imposed on ASMCs through lineage-specific differentiation leaves ascending-ASMCs mechanically maladapted in MFS.

**Figure 3:**
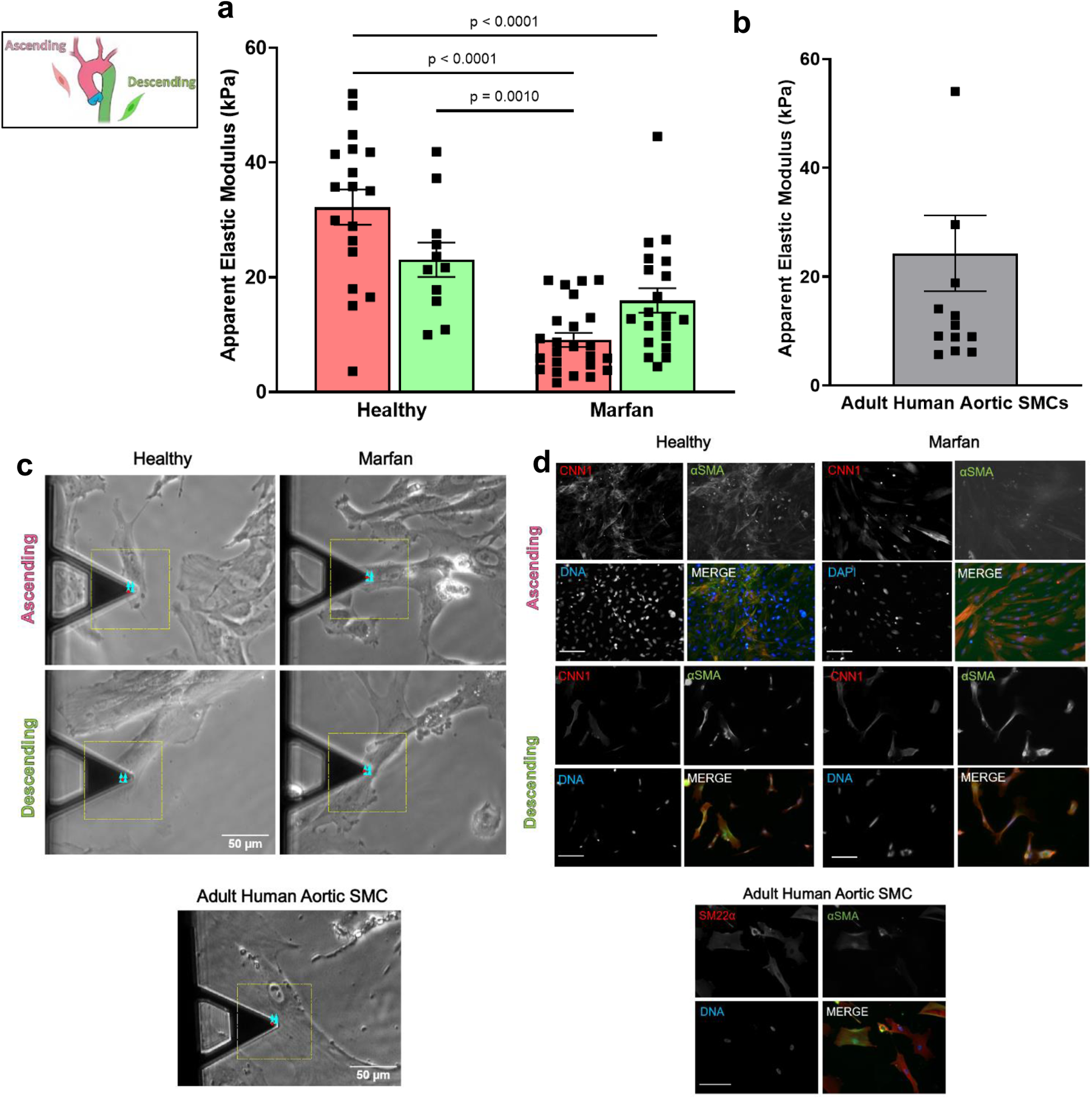
AFM reveals late-stage ascending ASMCs are uniquely less stiff in Marfan Syndrome. (a) AFM measured cell elastic modulus of each hiPSC-ASMC groups at a late cell-stage; Healthy ascending, healthy descending, Marfan ascending, Marfan descending. Ascending (neural crest derived) ASMCs are shown in pink while descending (paraxial mesoderm derived) are shown in green. Bars represent mean±SEM. Individual cells are shown as points (healthy: MSN09 & MSN22, Marfan: MFS44 & MFS60). p-values were calculated using an unbalanced multi-way ANOVA with Tukey-Kramer post hoc. (b) AFM measured cell elastic modulus of healthy primary adult human aortic SMCs. Bars represent mean±SEM. Individual cells are shown as points. (c) representative phase-contrast images of iPSC-ASMCs and primary ASMCs being measured via AFM. (d) Representative immunofluorescence images of fixed late-stage iPSC-ASMCs and healthy primary adult human aortic SMCs, scale bar = 200 μm.

### Marfan ASMCs have deficient calcium signaling

To test for functional deficiencies in late-stage ASMCs, we measured intracellular calcium response to 2.5 mM of the cholinergic agent carbachol, based on fluorescence intensity of the calcium sensitive dye, Fluo4AM (**Fig. 4**). The average response of both ascending and descending healthy ASMCs (n=9 cells/group) was a strong initial release signal, followed by robust cyclic waves of calcium cycling that decayed back towards basal levels over the course of several minutes (**Fig. 4a**). On the other hand, the average response from Marfan ASMCs (both ascending (n=11) and descending (n=16)) was an initial carbachol-induced calcium release, followed by a quicker decay to basal levels without propagation of robust waves (**Fig. 4c**). The MFS ASMC response was similar to the response of fibroblasts (**Fig. 4c**). In line with our data, similar functional abnormalities were previously seen in reduced contraction response of mesenteric arteries of the *Fbn1*^*C1039G/+*^ mouse MFS model ^41,42^. Together these observations indicate that Marfan iPSC-ASMC show functional changes in calcium signaling representative of those observed in the aortas of other MFS models ^43,44^.

**Figure 4:**
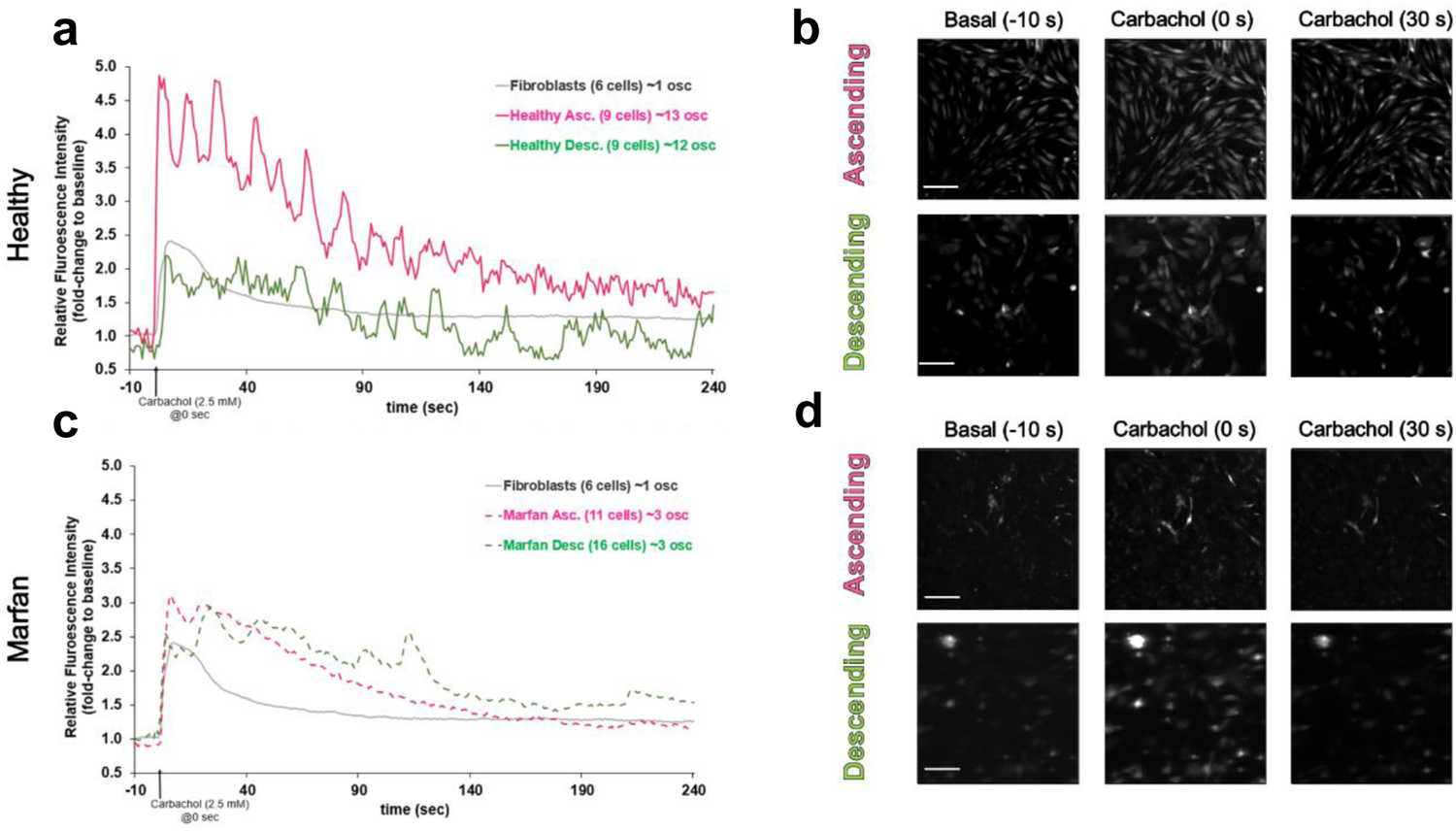
Marfan ASMCs are deficient at signaling calcium. (a) Healthy and (c) Marfan, late-stage ASMC fluorescence response to carbachol activation (2.5 mM added at time zero). Ca^2+^ flux was measured via the calcium sensitive dye, Fluo4AM, and quantified per cell relative to basal signal. Average number of contractions/oscillations (osc) for healthy, ascending (13 osc, n=9 cells) and descending (12 osc, n=9 cells). Average number of contractions/oscillations (osc) for Marfan, ascending (3 osc, n=11 cells) and descending (3 osc, n=16 cells). Each graph shows human dermal fibroblasts response for reference. (b,d) Example images of healthy and Marfan cell fluorescent response before, during, and after carbachol activation, scale bar = 200 μm.

### MFS Mechanical Phenotype Emerges at an Early Stage of ASMC Maturation

As hypothesized, in vitro models using late-stage matured hiPSC-ASMCs showed significant mechanical deficiencies in MFS vs. healthy groups, particularly for the neural crest lineage differentiations representing cells of the ascending aorta. Granata and colleagues similarly showed that their comparable hiPSC-ASMC S30 time-point models revealed significant differences in MFS SMC genes and phosphoproteins, but they were unable to see differences at their early post-SMC-differentiation timepoint ^12^. **Figure 5a** reveals that the mechanical phenotype seen in late-stage ASMCs (**Fig 3)**, is also seen at an early ASMC stage of post-differentiation maturation. Early-stage ASMCs were positive for ASMC markers (**Fig. 5c**) and exhibited mechanics similar to late-stage cells; where Marfan ascending ASMCs were significantly less stiff than descending (p=0.0209) (**Fig. 5a**). Most importantly, ascending ASMCs were significantly least stiff in MFS compared to healthy (p=0.0020) (**Fig. 5a**). This indicates that the MFS phenotype of intrinsically decreased compliance of ascending ASMCs is present at an early stage of cell maturation, and persists to a late maturation cell-stage.

**Figure 5:**
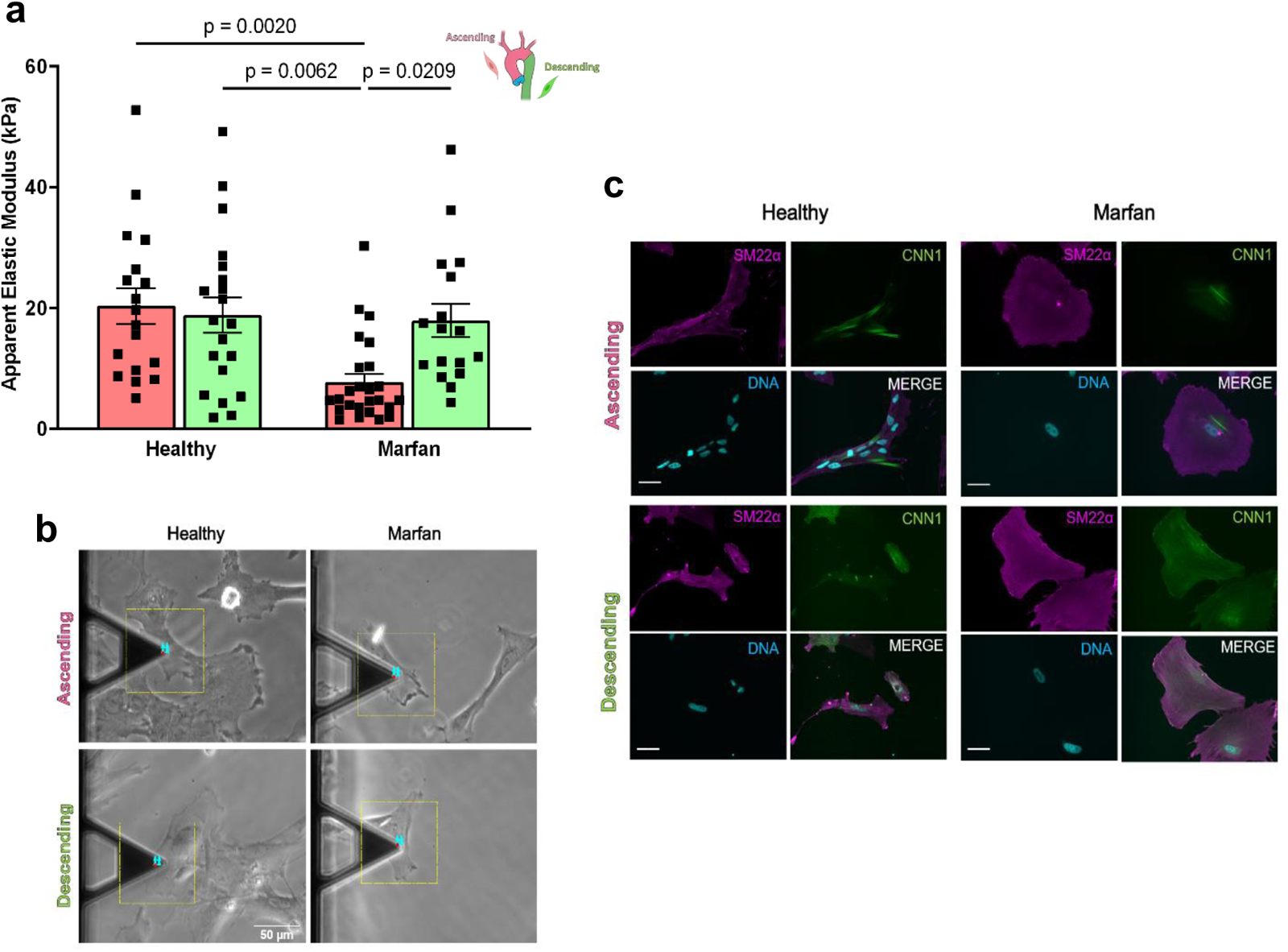
Early-stage Ascending ASMCs are less stiff in Marfan Syndrome. (a) AFM measured cell stiffness of each hiPSC-ASMC group at an early (day 6) post-differentiation maturation stage. Bars represent mean±SEM. Individual cells are shown as points (healthy: MSN09 & MSN22, Marfan: MFS44 & MFS60). p-values were calculated using an unbalanced multi-way ANOVA with Tukey-Kramer post hoc. (b) representative phase-contrast images of iPSC-ASMCs and primary ASMCs being measured via AFM indentation. (c) Representative immunofluorescence images of fixed early-stage iPSC-ASMCs, scale bar = 50 μm. SM22α; transgelin (magenta), CNN1; calponin (green).

## Discussion

This study employs an established MFS patient-derived hiPSC (human induced pluripotent stem cell) ASMC model, that has proven to faithfully mimic key aspects of the human vascular phenotype ^12^. Using this model, we have shown a connection between vascular SMC mechanics, ASMC developmental origin, and the aortic propensity of ascending aneurysm in Marfan syndrome. Smooth muscle cell stiffness has proven to be a promising biomarker of aortic pathology ^29,40,45,46^, that can translate across multiple biological scales from cells to tissues and whole organisms ^29,30^. Here we found that Marfan Syndrome offers a clinically important context to translate such metrics towards clarifying the evasive mechanisms driving ascending aortic aneurysm.

In this study a key finding was that neural crest lineage hiPSC-ASMCs representing cells of the ascending aorta, which would reside in the most aneurysm prone area of the Marfan aorta, were intrinsically less stiff, both at a late (**Fig. 3**) and early stage of cell maturation (**Fig. 5**) compared to healthy controls as well as diseased cells of the paraxial mesoderm lineage descending aorta. Importantly, Granata and colleagues found that in comparison to mesodermal hiPSC-SMCs, ectodermal hiPSC-SMCs in MFS had the most severely depleted fibrillin-1 phenotype ^12^. This phenotype was linked to neural crest (NC) specific deficiencies in their late-stage MFS SMCs. Particularly, they demonstrated MFS NC-SMCs had increased Itgb1, KLF4, and p38 which contributed to a stretch sensitive increase in cell death ^12^. If the aortic regions subject to the highest heamodynamic loads in vivo were also the weakest, it would reason that they are most vulnerable to aneurysm, but there is still debate over the heterogeneity and source of weakening. Mouse models, and to some extent clinical data, have shown that medial matrix degeneration occurs throughout the entire aorta in MFS, yet that alone is insufficient to guarantee aneurysm. Additionally, if the propensity for ascending aortic aneurysm in MFS was driven primarily by heamodynamic pressures, then lowering blood pressure to descending aortic levels should ameliorate aneurysm formation in the ascending aorta, which it does not ^19^. Researchers have noted that while both proximal and distal aortic SMCs appear indistinguishable at baseline, their developmental origin can imbue distinct responsiveness to cytokines. Avian models have shown that despite the different primary embryological origins of ectodermal and mesodermal derived SMCs, they expressed the same levels of differentiated ASMC markers in vitro ^47^. In contrast, neural crest derived ASMCs had dramatically increased DNA synthesis in response to TGFβ, when compared to mesodermal ASMCs ^47,48^. Furthermore, Owens and colleagues discovered in the ex vivo mouse aorta, that AT1aR mediated Angiotensin-II aortic thickening occurred via medial hyperplasia in the ascending aorta, while it did not in other aortic regions ^49^. With the advent of more human relevant models, Cheung and colleagues corroborated these findings in a hiPSC model, where they showed that MLK2 (a transcriptional co-activator of serum response factor), an embryogenesis driver in vivo, was also critical for in vitro SMC differentiation of neural crest cells, but not mesodermal cells ^11^. Moreover, their hiPSC model demonstrated that neural crest SMCs significantly proliferated in response to Angiotensin-II and TGFβ1, while mesodermal SMCs did not ^11^. Taken together this supports an emerging explanation of disease pathology, that active SMCs within the aortic media drive regional degradation in Marfan Syndrome, determined by developmental origin-based differences in their mechanobiology. Intimately linked to cell mechanics is cell function. We showed that compared to healthy, late-stage MFS ASMCs were less stiff. As well as functionally demonstrated that neural crest ASMCs in MFS have impaired calcium signaling (**Fig. 4**), aligned with other human and mouse studies ^12,41–44^. Studies showed that Marfan hiPSC-ASMCs have abnormal mechanosensing apparent through integrin presentation, cell adhesion, and tissue remodeling proteinases ^12^.

TGFβ, which is strongly implicated in MFS aortic aneurysm progression ^50,51^, is known to have robust embryogenic ^52,53^ and widespread biological effects across species ^54–56^. While the critical role of disrupted TGFβ signaling in MFS is well accepted, its temporally-dynamic effects, and thus the optimal timing of prophylactic therapeutic intervention, is uncertain. A group led by Dr. Francesco Ramirez showed in the *Fbn1*^*mgR/mgR*^ mouse that TGFβ has dimorphic effects on aneurysm development ^16^. Whether TGFβ neutralization was initiated before or after aneurysm formation determined if it exacerbated or mitigated aneurysm formation, respectively ^16^. While the *Fbn1*^*C1039G/+*^ MFS mouse showed that TGFβ-mediated Erk1/2 activation drives aneurysm formation ^51^, MFS hiPSC-ASMCs showed that by reducing TGFβ activity via Losartan, apoptotic activity was reduced in MFS SMCs, but not all aberrant mechanobiological pathways were mitigated. One possible explanation is the disconnect between rodent and human models. Notable are the angiotensin (AT) receptors; while humans express a single autosomal AT_1_ receptor gene, rodents express two related AT_1A_ and AT_1B_ receptor genes^57^. This was reflected in lackluster randomized clinical trials with MFS patients, where Losartan disappointingly showed no added benefit over placebo or beta-blockade ^19^. Interestingly the youngest patients appeared to receive the greatest therapeutic effect ^19,58–60^. As such we examined the mechanical phenotype and quantified protein levels of our early maturation-stage hiPSC-ASMCs. We saw that early-stage MFS hiPSC-ASMCs displayed the same intrinsic and matrix-independent weakening as our late-stage ASMCs, particularly with the ascending group being the least stiff in MFS (**Fig. 5**). While the study from Dr. Sinha’s group did not detect any differences in SMC RNAs at their early-stage of differentiation ^12^, our functional phenotype was present. To examine the potential disconnect between expressed RNA levels and protein abundances, we validated our early-stage phenotype findings via mass-spectroscopy quantification of cellular proteins. This revealed that while control proteins were unchanged (ACTB and TUBB1), cytoskeletal (ACTA2, CNN1, TAGLN) and contractile (MYH11, MYL6B) proteins were significantly depleted in MFS ascending ASMCs (**Figure 2**). Our data suggest intrinsic cell mechanics may offer an early biomarker in MFS, specific to ascending ASMCs. This validated biomechanical phenotype, which emerges at an early ASMC stage and persists into late-stage maturation, offers a novel biophysical marker of MFS pathology. Interestingly, we identified two significantly depleted proteins; integrin beta 3 (ITGB3) which acts with integrin alpha 5 to form a cell adhesion anchor to fibrillin-1 ^61^, and actinin alpha 1 (ACTN1) which is an actin cross linker that can reduce cell stiffness by destabilizing the actin cytoskeleton, and was shown to be a driver of abdominal aortic aneurysm in mice ^62^. Evaluation of the entire significantly depleted protein set revelated significant enrichment with Marfan-related processes (e.g. matrix organization, integrin adhesion, aortic aneurysm and dissection). While the reported protein targets were chosen partially based on their ability to explain our cellular phenotype, the dataset as a whole suggest its proteins are significantly associated with the clinical phenotype seen in Marfan Syndrome. Further exploration of the proteomic dataset, and validation experiments, are needed to understand how uniquely abundant and depleted proteins in ascending ASMCs drive the Marfan phenotype of ascending aortic aneurysm. Particularly, a finer-tuned study which considers cell adhesion to specific substrate coatings and their subsequent elicitation of integrin expression and impact on cell mechanics. This study is important because Marfan matrix is very different than healthy. If NC cells adhere and remodel differently than PM in MFS, this could explain the regional differences in aortic wall degeneration.

Within the research of ascending aortic aneurysm and Stanford Type A dissection in MFS, the question of aortic root versus ascending aorta comes into play. Contrasting hypotheses about the primary driver ASMCs are complicated by obscured localization of clinical aneurysm initiation and subsequent progression. It is important to note that in the aortic media; lateral plate mesodermal derived SMCs occupy the aortic root and pulmonary artery, neural ectoderm SMCs occupy the ascending aorta, and paraxial mesoderm SMCs occupy the descending/abdominal aorta. The medial SMC transition between ascending and descending aorta is a relatively sharp boundary, while the medial SMC change between root and ascending aorta is a more graded and intermingled transition ^5,9^. Thus, while the degraded region in aneurysm is apparent, the intermixed boundaries between aortic root and ascending aorta can veil its anatomical and developmental origin. Although neural crest hiPSC-SMCs were reported to have the most significant and comprehensive abnormalities in MFS, and not lateral plate mesoderm hiPSC-SMCs ^11,12^, which would support the uncommon clinical observation of pulmonary aortic dissection in MFS, others have found defective adhesion and SMC phenotype modulation unique to lateral plate mesoderm SMCs in similar models ^23,63–65^. One understudied mechanism that may play a role is the paracrine signaling between medial ASMCs within the root-to-ascending aorta transition. While it has been postulated that paracrine effects should be considered from an endothelial perspective ^66^, it is important to recognize that mechanical differences are predominantly observed in the aortic media layer ^13^. Further work is needed to dissect the differences in origin-based SMC mechanobiology and the connection with respective aortic medial boundaries.

## Conclusion

In conclusion, this study provides support for a paradigm shifting explanation that mechanobiological anomalies in neural crest derived smooth muscle cells of the ascending aorta drive the regional aneurysm propensity in MFS. While a picture is starting to form with SMC embryological origin at the center, the exact mechanisms defining the mechanobiology of aneurysm bias remain unclear. Detailing these pathways warrants more work. This research offers support for an ontogenic predisposition to ascending aortic aneurysm in MFS, and an early-stage physical biomarker of MFS pathology that can help guide research towards improved therapeutic targets for the most common and life-threatening manifestation of MFS.

## Acknowledgments

We want to thank; Christoph Schaniel (Department of Medicine, Division of Hematology and Medical Oncology) at the Icahn School of Medicine at Mount Sinai New York for generously donating healthy iPSC lines for study; The lab of Dr. Evren Azeloglu (Department of Pharmacological Sciences and Department of Medicine/Division of Nephrology) at the Icahn School of Medicine at Mount Sinai New York for offering guidance in conducting proteomic experiments; and the Rutgers University Proteomic Facility (Tong Liu, and Hong Li) for running our proteomic samples.

